# Occasional paternal inheritance of the germline-restricted chromosome in songbirds

**DOI:** 10.1101/2021.01.28.428604

**Authors:** Yifan Pei, Wolfgang Forstmeier, Francisco J. Ruiz-Ruano, Jakob C. Mueller, Josefa Cabrero, Juan Pedro. M. Camacho, Juan D. Alché, Andre Franke, Marc Hoeppner, Stefan Börno, Ivana Gessara, Moritz Hertel, Kim Teltscher, Ulrich Knief, Alexander Suh, Bart Kempenaers

**Affiliations:** Department of Behavioural Ecology and Evolutionary Genetics, Max Planck Institute for Ornithology, 82319 Seewiesen, Germany; Department of Organismal Biology, Evolutionary Biology Centre (EBC), Science for Life Laboratory, Uppsala University, SE-752 36, Uppsala, Sweden; School of Biological Sciences, University of East Anglia, Norwich NR4 7TU, UK; Department of Genetics, University of Granada, E-18071, Granada, Spain; Department of Biochemistry, Cell & Molecular Biology of Plants, Estación Experimental del Zaidín, Spanish National Research Council (CSIC), Profesor Albareda 1, E-18008 Granada, Spain; Institute of Clinical Molecular Biology, Christian-Albrechts-University of Kiel, Kiel, Germany; Sequencing Core Facility, Max Planck Institute for Molecular Genetics, Ihnestraße 63-73, 14195 Berlin, Germany; Department of Behavioural Neurobiology, Max Planck Institute for Ornithology, 82319 Seewiesen, Germany; Division of Evolutionary Biology, Faculty of Biology, Ludwig Maximilian University of Munich, Großhaderner Straße 2, D-82152 Planegg-Martinsried, Germany

**Keywords:** germline-restricted chromosome, paternal spillover, maternal inheritance, elimination efficiency, spermatogenesis, genetic conflict, selfish DNA, zebra finch, songbirds

## Abstract

All songbirds have one special accessory chromosome^1–4^, the so-called germline-restricted chromosome (GRC)^4–7^, which is only present in germline cells and absent from all somatic tissues. Earlier work on the zebra finch (*Taeniopygia guttata castanotis*) showed that the GRC is inherited only through the female line^4,6,8,9^ – like mitochondrial DNA^7,9–12^ – and is eliminated from the sperm during spermatogenesis^5,7,9–11^. Here we show that the GRC can also be paternally inherited. Confocal microscopy using GRC-specific FISH probes indicated that a considerable fraction of sperm heads (1-19%) in zebra finch ejaculates still contained the GRC. In line with these cytogenetic data, sequencing of ejaculates revealed that individual males from two families differed strongly and consistently in the number of GRCs in their ejaculates. Examining a captive-bred population of hybrids of the two zebra finch subspecies (*T. g. guttata* and *T. g. castanotis*) revealed that the descendants inherited their mitochondria from a *castanotis* mother but their GRC from a *guttata* father. Moreover, GRC haplotypes across nine different *castanotis* matrilines showed at best a weak tendency to be co-inherited with mtDNA haplotypes. Within *castanotis*, the GRC showed little variability, while the mtDNA of matrilines was highly divergent. This suggests that a single GRC haplotype has recently spread across the entire *castanotis* population, crossing the matriline boundaries via paternal spillover. Our findings raise the possibility that certain GRC haplotypes could selfishly spread through the population, via additional paternal transmission, thereby outcompeting other GRC haplotypes that were limited to strict maternal inheritance, even if this was partly detrimental to organismal fitness.

## Main

In sexually reproducing eukaryotes, the stable inheritance of the nuclear DNA typically requires recombination and segregation of pairs of homologous chromosomes that come from both parents. The songbird germline-restricted chromosome, GRC, is an intriguing exception^1,9,13^. As the name indicates, the GRC is only found in cells of the germline in all songbirds examined to date, but is absent from any somatic tissue^4,6,8,9^, presumably due to its elimination from somatic cells during early embryogenesis. While the functional significance of the GRC is still largely unknown^8^, this special chromosome appears to be far more than just an accumulation of highly repetitive DNA, as it may initially have appeared^12^. To the contrary, the zebra finch *Taeniopygia guttata castanotis* GRC is rich in genes that are expressed in testes or ovaries^8^, and the gene content of the GRC appears to be evolving rapidly^8,14^ leading to remarkable variation in GRC size between species^4,5^. This rapid evolution is puzzling, because the GRC’s adaptive value for passerines is not at all obvious (compared to all other birds that lack a GRC^4^). Rapid evolution often takes place when some genetic elements successfully manipulate their mode of inheritance to their own advantage (so called ‘selfish genetic elements’^15–17^). Hence, as a key step towards understanding the evolution and the function of the GRC, it appears essential to fully understand how the GRC is inherited.

In most studies to date, the GRC was observed in two copies in female (primary) oocytes^4,5,7,12^, and as a single chromosome in male spermatogonia^4–9^. Cytogenetic investigations in males (predominantly *T. g. castanotis*) found that this single-copy GRC is eliminated from nuclei during meiosis I and expelled from spermatids in late spermatogenesis^5–7,9–12^. Based on observations from multiple species^5–7^, it has been concluded that the avian GRC is exclusively maternally inherited. Yet to date, the mode of inheritance of the GRC remains untested genetically, implying that suggestions about the evolutionary significance of the GRC^5,8,13^ remain speculative. In this study, we used the two zebra finch subspecies *T. g. castanotis* of Australia (hereafter *castanotis*) and *T. g. guttata* of the Indonesian islands such as Timor (hereafter *guttata*) to address the issue of inheritance. Specifically, we combined cytogenetic and genomic data to study (a) the elimination efficiency of the GRC during spermatogenesis, (b) the strictness of the proposed matrilineal inheritance (i.e. expected co-inheritance with the mitochondrial genome) and (c) the genetic variation of the GRC within the *castanotis* subspecies.

## GRC-specific sequences in ejaculates: repeatability and differences between families

In principle, GRC-specific sequences might be found in ejaculates in three different forms: (1) expelled free-floating GRC micronuclei^9,11^, (2) small, digested DNA fragments^10,11^ and (3) non-expelled GRCs or parts thereof in sperm heads. According to current knowledge^4–7^, primary spermatocytes contain a single copy of the GRC that is being expelled as a micronucleus during early meiosis. As each primary spermatocyte results in four haploid spermatozoa and all chromosomes have two chromatids, we expect that ejaculates contain up to 25 free-floating GRC-micronuclei per 100 spermatozoa, in case of 100% expulsion. We examined 7 natural ejaculate samples from 5 different *castanotis* males, using a probe for the GRC-linked high copy number gene *dph6*^8^ for fluorescent *in situ* hybridization (FISH) to label the GRC in the ejaculate. Contrary to the expectation, we frequently found the FISH signal for the GRC inside sperm heads (mean = 9% of the sperm heads, range: 1-19%) and only a few free-floating micronuclei (mean = 1 micronucleus per hundred sperm heads, range: 0-2; **Fig. 1**; **Supplementary Table 1**). The GRC-containing spermatozoa (*dph6*-positive) showed no visible morphological differences to the GRC-negative ones (**Fig. 1**). A subsequent confocal microscopy analysis of the *dph6*-positive sperm heads showed that the signal came from inside the sperm nucleus (**Fig. 1c**; **Supplementary video**). These results show that, in the zebra finch, the GRC is not completely eliminated during spermatogenesis, and imply that a non-negligible number of spermatozoa can potentially transmit the GRC.

**Fig 1.**
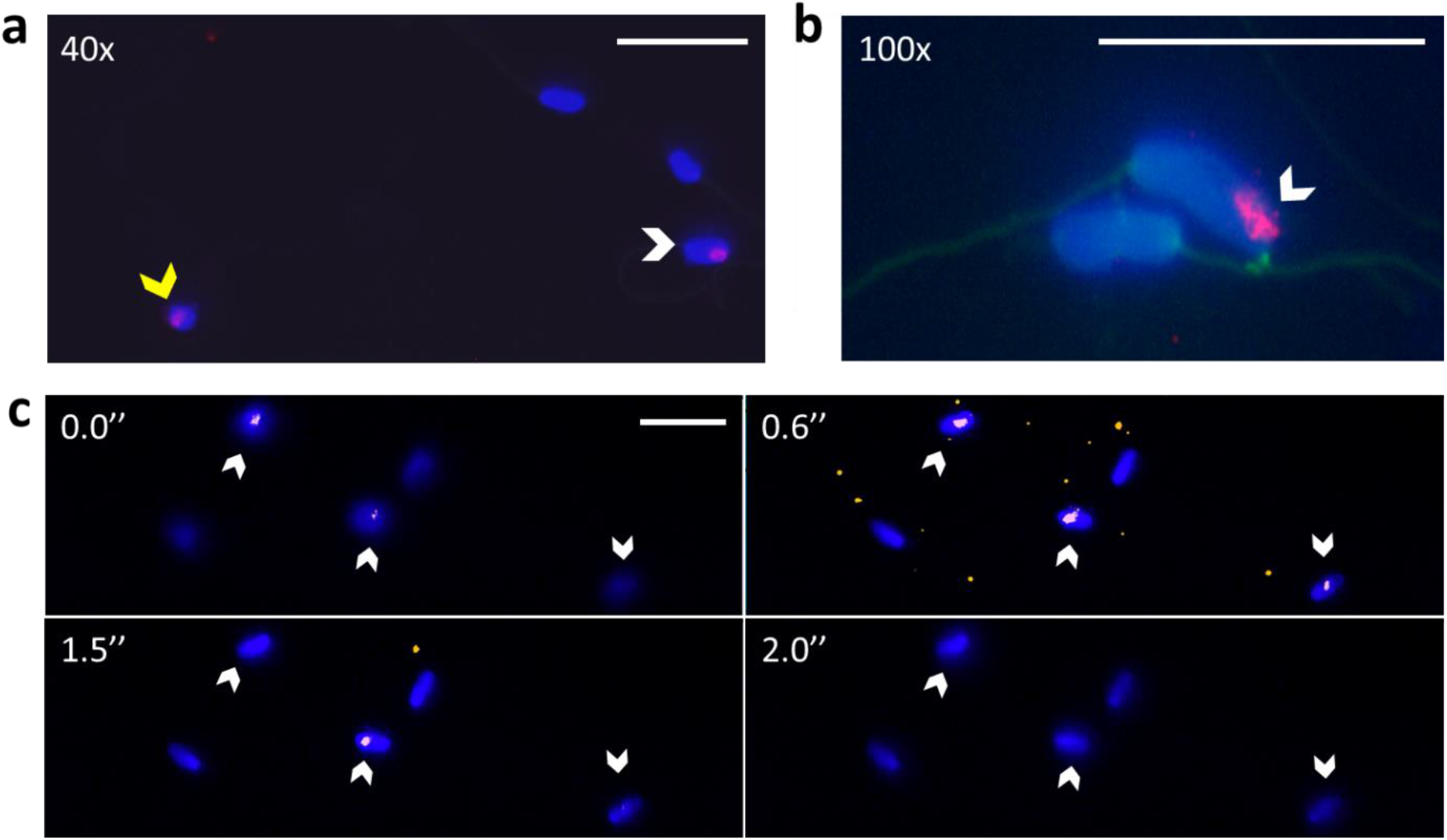
Cytogenetic evidence for the presence of the GRC in the nucleus of zebra finch *Taeniopygia guttata castanotis* sperm. The GRC-amplified probe *dph6* (see Methods) indicates the presence of the GRC (red) inside some sperm heads (white arrow in **a**-**c**) and in free-floating micronuclei (yellow arrow in **a**). Blue DAPI stain without red indicates sperm heads without GRC. Green autofluorescence shows the sperm flagellum (in **b**). **a**, 40X magnification. **b**, 100X magnification. **c**, Individual z-sections under a confocal microscope show the sequential appearance and disappearance of the *dph6* signal along consecutive sections, indicating the location of the GRC within the nucleus of the spermatozoa. Time (in seconds) refers to the **Supplementary video.** The video consisted of 24 sections representing a total of 6.0 μm depth. Scale bars are 20 μm.

Given the observed variability in the proportion of sperm heads that contained the GRC, we estimated individual repeatability and between-family variation in the proportion of sperm heads that contained the GRC. To do this, we quantified the amount of GRC in ejaculates using PCR-free Illumina sequencing and compared the results to those obtained from the cytogenetic data (presented above). As GRC-linked sequences are often difficult to distinguish from their ancestral A-chromosomal paralogs (as the latter vary substantially between individuals)^8^, GRC content is most easily quantified by focussing on sequences that reside on the GRC in numerous copies (compared to just two A-chromosomal copies). Hence, we first defined a set of high-copy-number sequences on the GRC. This was done by sequencing (for details see **Methods** and **Supplementary Table 2**) both testis (high GRC content) and liver (or muscle) samples (a GRC-free control tissue) of nine *castanotis* males from nine different matrilines (i.e. mitochondrial haplotypes) that represent most of the genetic variation of the mitochondria in the wild (**Extended Data Fig. 1**). Because there is no GRC reference assembly available, we only looked at GRC-linked regions that have an A-chromosomal paralog^8^. Hence, we mapped each of the nine testis and nine liver (or muscle) libraries onto the soma-based reference genome taeGut1^18^, and then identified 1,742 windows of 1 kb length that were more than 4-fold enriched in coverage in all nine testis libraries compared to their respective liver library (i.e. log_2_ testis-to-soma coverage ratio larger than 2; **Fig. 2a**).

**Fig 2.**
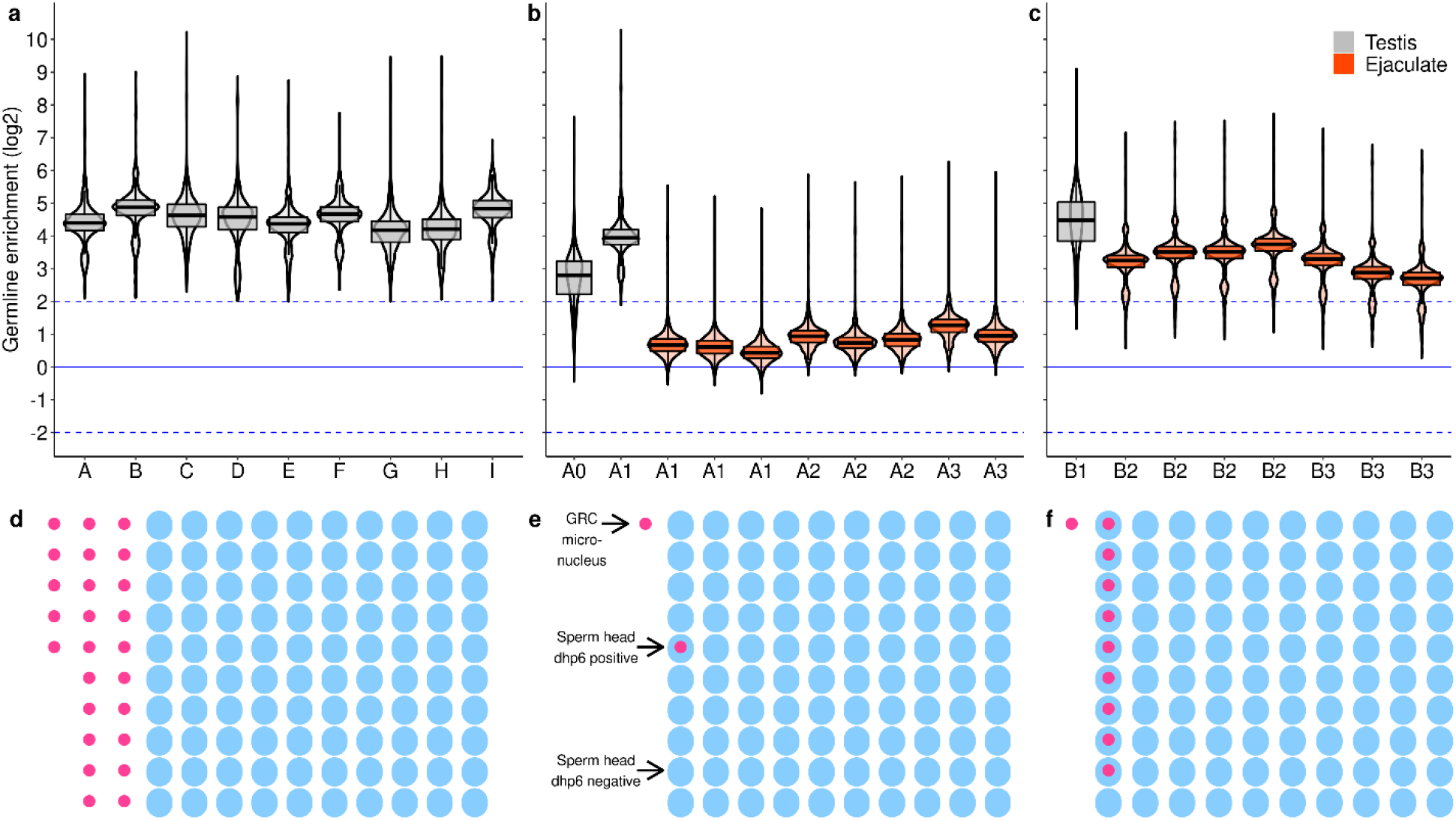
The elimination efficiency of the GRC differs between *castanotis* matrilines A and B. **a-c,** Comparison of sequencing coverage of GRC-containing (testis, indicated in grey, and ejaculate, orange) and GRC-free tissue (liver) identifies sequences that are GRC-linked in high copy number^8^. The solid blue line refers to a log_2_ germline-to-soma coverage ratio = 0, i.e. no germline enrichment; the dashed blue line refers to a 4-fold increase (top) of sequencing coverage in germline compared to soma tissue. **a,** Violin-box plots show coverage ratios of the selected sequences (N = 1,742 windows of 1 kb) with more than 4-fold (log_2_ > 2) enrichment in testes in comparison to soma in all nine *castanotis* males (A to I, of different matrilines). In boxes, thick horizontal lines show the median, and boxes indicate the 75th and 25th percentiles, respectively. **b**, Coverage ratios of the selected sequence windows in matriline A (four related individuals A0-A3) show that ejaculates contain lower amounts of GRC-derived reads compared to testes (comparison of the median of eight ejaculates with two testis samples: b = −2.56, SE = 0.29, P < 0.001; **Supplementary Table 3**), as expected from previous work^10,11^. **c,** Coverage ratios of the selected sequences in matriline B (three related individuals B1-B3) with a higher GRC content in ejaculates and hence a smaller difference with testis (comparison of the median of seven ejaculates with one testis sample: b = −1.21, SE = 0.39, p = 0.02; **Supplementary Table 3**). **d-f,** illustration of the expected (**d**) and observed (**e**, **f**) number of sperm heads that are free of GRC (blue ovals) and that contain GRC (blue ovals with a red circle inside; i.e. positive *dph6* signal, see Fig. 1), as well as the number of free-floating GRC-micronuclei (red circles). **d,** Error-free expulsion of GRC from spermatocytes (expected based on previous work) should result in up to 25 free-floating GRC micro-nuclei per 100 sperm heads in the ejaculate^6,7,9,11^. **e,** Ejaculates from matriline A showed 1% of GRC-positive sperm heads (N = 677 scored sperm). **f,** Ejaculates from matriline B showed 9% of GRC-positive sperm heads (N = 1,533 scored sperm).

After identifying these GRC-linked windows, we examined the GRC content of ejaculates. We re-sequenced 15 ejaculates (2-4 samples per male) together with blood samples of their parents (as GRC-free control tissue) from five additional *castanotis* males stemming from two families using PCR-free Illumina technology. These five *castanotis* males were from the matrilines A (N = 3) and B (N = 2), as described in ^19^, and include one male per family for which we had obtained cytogenetic data (as presented above, see **Supplementary Table 2**). The two matrilines represent the two most common mitochondrial haplotypes in the domesticated zebra finch (*castanotis*) ‘Seewiesen’ population^20^ (see **Extended Data Fig. 2**). As GRC-positive controls for the ejaculate samples, we also sequenced three testis samples (and the respective liver samples as GRC-free controls) from families A and B (PCR-free Illumina technology; **Supplementary Table 2**). After mapping the above sequenced libraries, we estimated the log_2_ germline-to-soma coverage ratio of the 1,742 selected windows (for the 15 ejaculates and three testes from six brothers and one uncle of matrilines A and B compared to their corresponding soma samples).

Based on coverage ratios, ejaculates from the same individual male zebra finch were remarkably repeatable in their GRC content (**Fig. 2b-c**; R_male_ = 0.82, P < 0.001; **Supplementary Table 3**). The majority of the repeatable variation was due to a between-matriline difference (R_matriline_ = 0.86, P < 0.001; **Supplementary Table 3**). Ejaculates from males of matriline B had significantly higher amounts of GRC (**Fig. 2b**) than those from males of matriline A (**Fig. 2c**;b_matrilineB_ = 1.7, SE = 0.21, P < 0.001; see also **Extended Data Fig. 3c-e,g-h, Extended Data Fig. 4** and **Supplementary Table 3**), confirming the results from the cytogenetic analysis (1% versus 9% of sperm heads were GRC-positive, respectively; **Fig. 2d,e**).

These results indicate that certain GRC haplotypes (e.g. those in matriline B) are more likely to be transmitted via sperm than others (e.g. those in matriline A), and hence more likely to potentially spread in a ‘selfish’ manner. The high individual repeatability and consistency within matriline raises the question whether the observed difference in the elimination efficiency of the GRC during spermatogenesis might have a relatively simple genetic or epigenetic basis. Future studies should test whether this is due to the GRC haplotype itself (including epigenetic marks such as histone or DNA modifications) or due to epistatic interactions between the GRC and the A-chromosomal genome.

## Paternal inheritance of the GRC in a hybrid population

Given the observed occurrence of the GRC in spermatozoa, we tested for paternal inheritance of the typically maternally-transmitted GRC^5,7,9,12^. First, we identified polymorphic markers that reliably distinguish different GRC haplotypes. Then, we looked for cases in which the maternally transmitted mitogenome and the GRC markers were not co-inherited (i.e. informative paternal transmission of the GRC). This search rests on the assumption of strict maternal inheritance of mtDNA haplotypes, which is highly likely because (1) paternal inheritance of the mitogenome is known to be rare^21^ and was usually detected in individuals with mitochondrial diseases and heteroplasmy^22,23^, and (2) the avian W chromosome and mitochondrial DNA are in high linkage disequilibrium^21^. We used data on all 12 *castanotis* males stemming from nine matrilines from which testes and soma samples had been sequenced (**Fig. 2a-c**), and we additionally sequenced a pair of testis and liver (**Supplementary Table 2**) of a *castanotis* x *guttata* hybrid descendant male with *guttata* phenotype but *castanotis*-introgressed genotype (i.e. *castanotis* matriline B and 5% of *castanotis* introgressed tracts on an otherwise *guttata* A-chromosomal genome), using 10X linked-read technology (**Fig. 3**; see **Supplementary material** for details).

**Fig 3.**
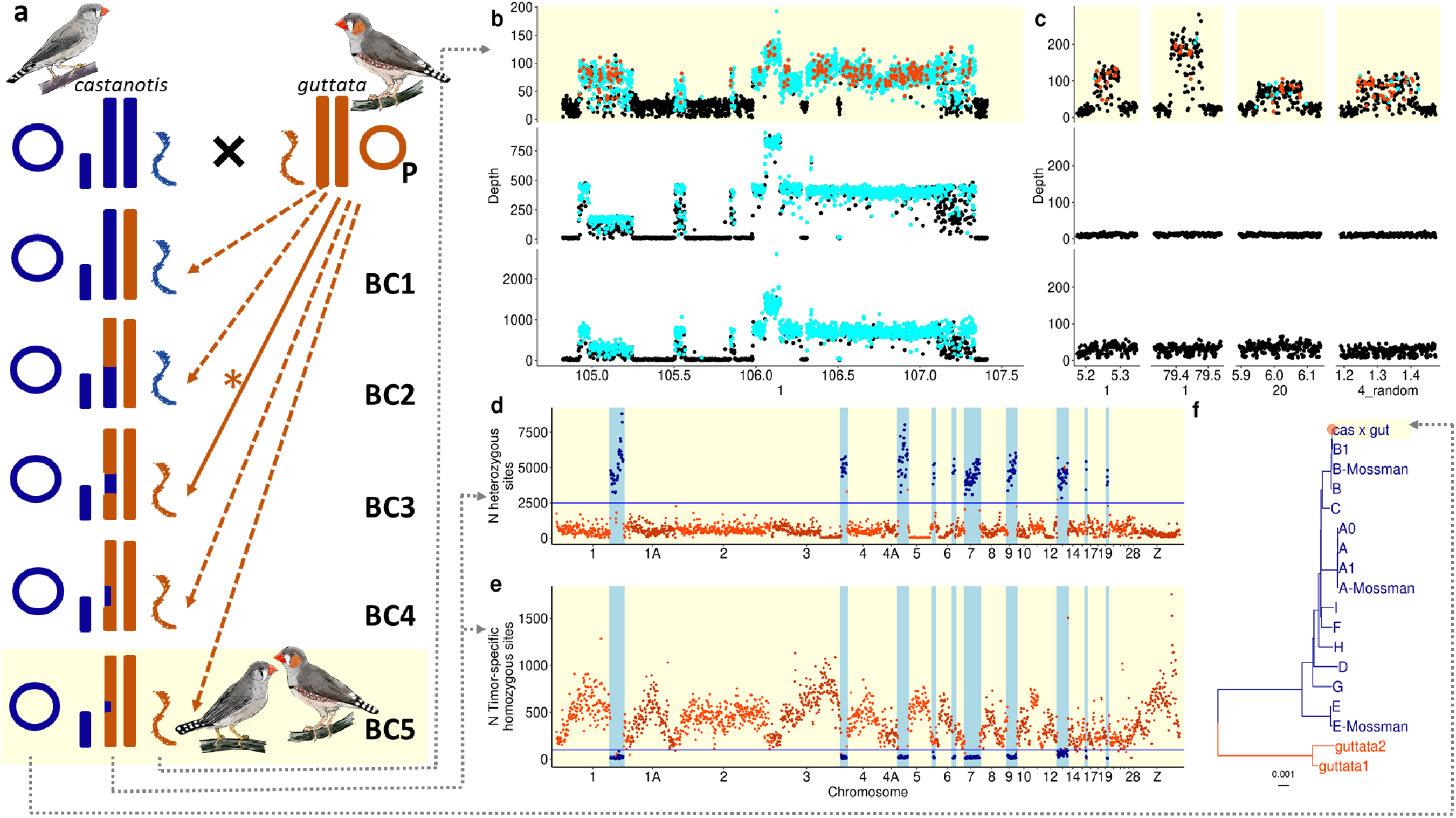
A case of paternal inheritance of the GRC in a captive-bred hybrid *castanotis* x *guttata* population. **a,** Reconstructed breeding history during domestication of the recently wild-derived *guttata* subspecies. Presumably because females of wild-derived *guttata* birds do not easily reproduce in captivity, we hypothesize that *guttata* males were crossed with an already domesticated *castanotis* female (from Europe), and the resulting female hybrids were back-crossed with pure *guttata* males for about five generations (P-BC5) until the population was phenotypically *guttata*-like. This reconstruction is based on the genotyping of one male^52^ of the resulting hybrid population (yellow background, bottom), which is characterized by a *castanotis* mother contributing the mtDNA (dark blue circle), the female-specific W-chromosome (dark blue short rectangle; see **Supplementary Results**) and 5% of the A-chromosomal DNA (blue fragments in the long rectangles), and *guttata* males contributing 95% of the A-chromosomal DNA (orange-brown long rectangles) and the GRC (orange-brown sigmoid symbol). Note that the paternal spillover of the GRC must have happened sometime between generations P and BC5 (solid arrow with asterisk). **b-c**, Raw read depth of selected genome regions of testis libraries (mapped to somatic-reference taeGut1^18^) for a *castanotis x guttata* hybrid (top row of each panel, yellow background) and two representative *castanotis* individuals (E and B; bottom two rows). GRC-specific sequences are distinguishable from somatic reference-like sequences by high read depth values and by testis-specific SNPs (cyan). Each dot represents a 1-kb window. Each cyan dot indicates a 1-kb window that contains ≥ 3 germline-specific SNPs. Each orange-brown dot shows a 1-kb window that contains private testis-specific SNPs. Note that only the hybrid has individual testis-specific SNPs in the GRC-linked sequences that are shared between all *castanotis* males and this hybrid individual (**b**) as well as private GRC-linked sequences that are absent in *castanotis* individuals (**c**), suggesting that it carries a *guttata* GRC. See **Extended Data Fig. 5** for genome-wide plots and more individuals. **d-e** Number of heterozygous sites (**d**) and number of fixed *guttata*-specific homozygous sites (**e**) in autosomal sequences including Z from somatic tissue of the *castanotis* x *guttata* hybrid compared to the sequences from a pooled DNA sample from 100 wild-caught *castanotis* zebra finches^27^. The horizontal blue line indicates the cut-off of 2500 heterozygous sites (**d**) or 100 fixed *guttata*-specific homozygous sites per 500-kb window (i.e. each dot; **e**). Dark blue dots show windows that contain an excess number of heterozygous sites (**d**), or a reduced number of *guttata*-specific homozygous sites (**e**), and indicate 10 introgressed *castanotis* segments. **f,** Phylogenetic tree of the mitogenomes, showing that the hybrid’s mtDNA (‘cas x gut’, orange-brown dot at the top of the tree) clusters with typical captive European *castanotis* zebra finches (dark blue) rather than with mitogenome assemblies of two published *guttata* datasets (SRA accession numbers SRR2299402^53^ and SRR3208120^54^ respectively; orange-brown). The scale bar shows the number of substitutions per site.

If the GRC would have been inherited exclusively through the matriline^5,7,12^, we expected that the hybrid male would show the same GRC haplotype as is typical for *castanotis* matriline B. However, we found a novel GRC haplotype that uniquely diverged from all the 12 GRC haplotypes examined so far (**Fig. 3b,c**; **Extended Data Fig. 5**). Given this high degree of divergence relative to the 12 *castanotis* GRCs, we hypothesized that this might be the hitherto unknown GRC of *guttata* (or at least a recombinant *guttata* x *castanotis* GRC). The high divergence was apparent in terms of (a) a high number of private testis-specific SNPs, i.e., SNPs that were only detected in the testis of this individual in GRC-amplified regions that were shared among all 13 GRCs (hence GRC-linked variants; N_high-confidence SNP_ = 312 in **Fig. 3b**; **Extended Data Fig. 5**), and (b) four independent regions that appear to be GRC-linked in high copy number in this individual but are absent from all other 12 *castanotis* GRCs (N_high-confidence SNP_ = 169 in **Fig. 3c; Extended Data Fig. 5;** including 13 genes with GRC paralogs that were not found previously^8^, see **Supplementary Table 4**; also see **Supplementary Table 5**). These results suggest that the GRC was inherited from at least one of the potential *guttata* males during back-crossing (**Fig. 3a**). We assume that the *guttata* GRC coexisted with, recombined with or replaced the *castanotis* GRC, and stably co-inherited with the *castanotis* mitochondria thereafter (**Fig. 3**).

## Reduced genetic diversity in low-copy-number genes on the GRC

We found surprisingly little variation among the 12 *castanotis* GRCs from nine matrilines, some of which were old matrilines as judged from mtDNA divergence (**Fig. 3f, Fig. 4a**, **Extended Data Fig. 1**). In sharp contrast to the highly diverged *guttata* GRC which harboured hundreds of private testis-specific SNPs (top row in **Fig. 3b-c** and **Extended Data Fig. 5a; Supplementary Table 5**), we did not find a single high-confidence GRC-specific SNP that was private to one of the 12 *castanotis* individuals (middle and bottom rows in **Fig. 3b-c** and **Extended Data Fig. 5b-m**).

**Fig 4.**
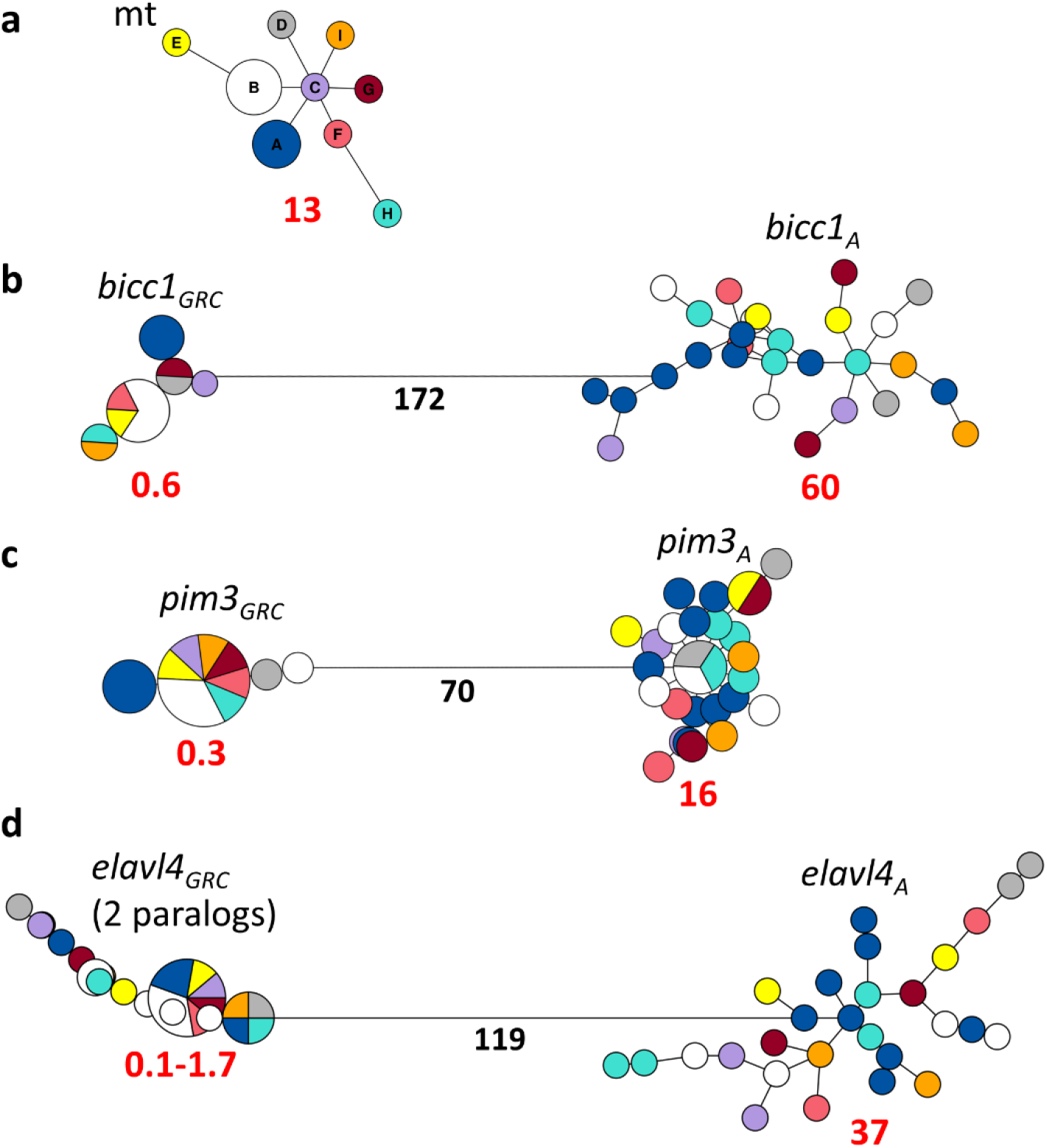
Haplotype networks showing the genetic diversity in GRC-linked genes (*gene*_GRC_) compared to their A-chromosomal paralogs (*gene*_A_) and the mtDNA (mt) in *castanotis* zebra finches. Shown are gap-free alignments of mitogenomes (**a**), two single-copy genes on the GRC^8^ (**b,**13,822 bp; **c,**3,286 bp) and a low-copy-number gene (presumably two paralogs) on the GRC^8^ (**d,**17,492 bp) with the two haplotypes of the respective single-copy A-chromosome paralogs from all *castanotis* germline samples used in this study (N = 14 individuals). Colors represent the different mitogenome haplotypes. The size of each circle indicates the number of individuals of each haplotype, and the length of the black lines corresponds to the number of mutational steps between haplotypes. Red numbers refer to the number of SNPs per kilobase for each cluster of haplotypes; black numbers refer to the number of mutations per kilobase between the GRC-linked and A-chromosomal paralogs (**b-d**). Note the highly reduced genetic diversity (red numbers) in the GRC genes (which corresponds to a mean number of pairwise differences per site π = 0.00015-0.00027; **Supplementary Table 6**) in comparison to their A-chromosomal paralogs (π = 0.0013-0.011; **Supplementary Table 6** and **Extended Data Fig. 7**; **b-d**). Further note that different mitogenome haplotypes may share the same GRC haplotypes (i.e. the circles contain multiple colors in GRC genes in **b-d**), indicating that mtDNA and GRC are not always inherited together.

We studied the within-population variation of the GRC (in *castanotis*), and tested for linkage disequilibrium between the GRC haplotypes and matrilines, by focusing on single- or low-copy-number GRC genes that were highly diverged from their A-chromosomal paralogs. The latter is necessary, because only high divergence ensures that all reads map without error to their correct origin, being either from the GRC or from the A-chromosomal paralog. We screened the published list of 267 GRC-linked genes^8^ for a high number (N > 50) and high density (> 3 per kb) of germline-specific SNPs, and identified four such genes (*pim3*, *elavl4*, *bicc*1 and *cpeb1*). The gene *cpeb1* was dropped because we were unable to assemble a GRC contig that was longer than two kilobases (see Methods). For the remaining three genes we assembled *de novo* a consensus contig that was longer than three kilobases using MEGAHIT v1.2.6^24^ (see Methods). We then constructed a GRC haplotype for each of the testis samples using the contrast with the somatic (liver or blood) samples of the 12 *castanotis* males, and for the pooled ejaculate samples of the 2 males from matriline B (to maximize the sample size; males B2 and B3 in **Fig. 2c**). Our assembled contigs of *bicc1*, *pim3* and *elavl4* were clearly GRC paralogs, because there were zero reads mapped onto them from the somatic libraries.

The GRC paralogs of *pim3* and *bicc1* were clearly in single copy, because all GRC mappings were homozygous (also see **Extended Data Fig. 6**). This confirms previous work showing that male germline cells carry a single copy of the GRC^4,5,7–9,12^. For the GRC paralog of *elavl4*, all germline samples shared the same heterozygous sites with allele frequency of ~50% (**Extended Data Fig. 6**), suggesting that the GRC contains two copies of *elavl4*. Therefore, we artificially phased two GRC haplotypes for each germline sample by keeping the reference allele for haplotype 1 while using the alternative allele for haplotype 2. This introduces a bias towards higher genetic diversity of haplotype 2, but this was not problematic for our purpose.

We compared the genetic diversity both among the GRC haplotypes, and between the GRC and their A-chromosomal paralogs using the three previously mentioned GRC-linked genes. For this, we aligned the haplotypes of the GRC (*gene_GRC_*) and their A-chromosomal paralogs (*gene_A_*) using MAFFT v7.429^25^, and then analysed the alignments using the R package pegas v0.14^26^. The A-chromosomal paralogs of the three genes represented the range of genome-wide autosomal genetic diversity found in *castanotis* in the wild^27^, with *pim3*_*A*_ representing low levels of diversity (16 SNPs per kb), *elavl4*_*A*_ representing average levels (37 SNPs/kb), and *bicc1*_*A*_ representing high levels (60 SNPs/kb) (**Fig. 4b-d right; Supplementary Table 6** and **Extended Data Fig. 7**). We also found considerable genetic diversity in mitochondrial DNA (13 SNPs/kb). In contrast, the two single-copy GRC-linked genes and *elavl4*_*GRC*_ showed low genetic diversity (*pim3_GRC_*: 0.3 SNPs/kb, *bicc1_GRC_*: 0.6 SNPs/kb, *elavel4*_*GRC*_ haplotype 1: 0.1 SNPs/kb and haplotype 2: 1.7 SNPs/kb; **Fig. 4**).

Given the low genetic diversity of the GRC, it is difficult to assess with confidence whether GRC haplotypes are co-inherited with the mitochondrial genome. For the two single-copy GRC genes, the *castanotis* males that have the same mitochondrial haplotype also have the same GRC haplotype (3 males of matriline A, and 4 males of matriline B; see **Extended Data Fig. 6**). Note, however, that most of these males are brothers that can be expected to share the same mitogenome and GRC-haplotype (only one male, B, is not closely related; **Supplementary Table 2**). Further, some males have different mitochondrial haplotypes, but share the same GRC haplotype (**Fig. 4b-d** and **Extended Data Fig. 6**). These observations, based on only 11 polymorphic sites on the two single-copy genes on the GRC (**Supplementary Table 6**), suggest that GRC and mtDNA are not always co-inherited in *castanotis* zebra finches.

## Discussion

Our results demonstrate that certain GRC haplotypes can be and have been transmitted from the father (**Fig. 3**) via sperm (**Fig. 1**), in addition to their regular maternal transmission. The likelihood of paternal inheritance of the GRC seems family-specific (**Fig. 2**) and may thus be heritable. Although some of our findings are based on a single case (**Fig. 3**) or on 15 ejaculates from two families (**Fig. 2**), taken together they clearly indicate that the GRC can be paternally inherited.

It remains unclear what happens when the two GRCs (i.e. one maternally and one paternally inherited) enter the embryo. However, we did not observe any GRC-heterozygous birds (**Extended Data Fig. 6**), suggesting that one GRC is removed (either always the same one, i.e. an efficient driver, or with a 50:50 probability). In either case, the GRC that escapes elimination during spermatogenesis has an evolutionary advantage by having two possible routes of inheritance (as opposed to a single route for other GRC haplotypes that are only maternally transmitted)^15^, which sets the stage for selfish evolution of DNA. The fact that we see low diversity on the *castanotis* GRCs even across highly diverged matrilines (**Fig. 4**), suggests that such a selfish GRC may have spread relatively recently. The highly diverged *guttata* GRC (**Fig. 3**) rules out the alternative explanation that the selected GRC genes have an unusually low mutation rate or experience strong purifying selection, which is also contradicted by the observed dynamic GRC evolution in general^4,8^.

Selfish evolution of the GRC might be widespread across songbirds, because we find evidence for paternal inheritance in both zebra finch subspecies and because of the observed similarity of meiotic behaviour (i.e. elimination during male spermatogenesis and recombination in female oocytes) of the GRC across all songbirds examined to date^5–7^. Interestingly, Malinovskaya et al.^5^ reported a GRC copy number mosaicism in spermatogonia and pachytene spermatocytes in males of the pale martin (*Riparia diluta*). We hypothesize that the spermatogonia and spermatocytes that possess an extra copy of the GRC might result in GRC-carrying sperm. The GRC-carrying sperm thereby brings the GRC a “selfish advantage”, meaning that such GRCs might be able to spread even if they were mildly deleterious to the organism (e.g. to a certain sex or developmental stage).

We observed a remarkable polymorphism in the efficiency of GRC-elimination from sperm and we expect that this will be mirrored in the ability of the GRC to spread paternally (**Fig. 2**). Such variation could only be evolutionary stable if the obtained advantages via selfish spreading would be compensated by other disadvantages, for example, if paternal inheritance would reduce fertility or embryo survival (antagonistic pleiotropy). A paternally-spreading GRC haplotype may also have been fixed in the population, as indicated by the low genetic diversity (**Fig. 4**) and the ability to spread in both zebra finch subspecies (**Fig. 2** and **Fig. 3**). After a certain GRC haplotype has successfully spread to the entire population (going to fixation), its ability to spread through this second (i.e. paternal transmission) route may lose its adaptive value, because there is no alternative haplotype to compete with. However, a second variant that lacks this paternal spreading ability could then only have invaded if it conveyed another advantage (e.g. to organismal fitness).

A polymorphism in the efficiency of elimination of the GRC during spermatogenesis (**Fig. 2)** might also explain why previous cytogenetic work on testes of songbirds failed to detect GRC-positive spermatozoa^4–6,8,9^. By chance, the examined individuals might resemble those in matriline A (**Fig. 2b**), in which the GRC elimination efficiency during spermatogenesis is high (99%).

Recent studies suggest a highly dynamic nature of the content of the GRC. Across species, there is excessive variation in size and content of the GRC^4,5^, especially when compared to the highly syntenic A chromosomes in birds^28–30^. In the Australian zebra finch (*T. g. castanotis*), much of the content of its GRC appears to have been derived from A-chromosomal paralogs only recently^8^. The zebra finch (*T. g. castanotis*) GRC is enriched for genes showing gonad-specific expression^8^, and some genes show signals of strong positive or purifying selection^8,14^, suggesting an essential role of the GRC in sexual reproduction. The genetic diversity of the three examined genes on the GRC is 40-fold lower than the diversity of their A-chromosomal paralogs (**Fig. 4**; **Supplementary Table 6**; also see **Extended Data Fig. 7** and ^31^). Other non-recombining sex-specific chromosomes also show highly reduced genetic diversity compared to their autosomes, possibly due to strong sexual selection (e.g. the human Y-chromosome shows a 23-41-fold reduction in the mean number of pairwise differences per site, i.e. π ^32^), or due to Hill-Robertson interference with the mitogenome (e.g. the avian W chromosome shows a 46-104-fold reduction of π in four *Ficedula* flycatcher species^21^- and a 90-fold reduction in chicken *Gallus gallus*^33^).

Another implication of the GRC’s strong linkage disequilibrium with the two other non-recombining elements in songbirds, i.e. the W-chromosome and the mitogenome, is that it reduces the efficiency of positive or negative selection on them, especially in small populations (i.e. Hill-Robertson interference)^34,35^. However, this is only true if these three elements are always co-inherited. With paternal spillover demonstrated here, the GRC is immediately decoupled from the other two, and can create a new optimal combination of the three elements – the GRC, the mitogenome and the W-chromosome – such that this new combination can rise in frequency. Thus, the gene richness and highly dynamic nature^4,8^ of the GRC and our observation that it can be paternally inherited offers novel opportunities to study genetic compatibility and the evolution of the W chromosome and the mitogenome^21^ of songbirds, which might be relevant for understanding the underlying genetic basis of speciation and the high species richness in songbirds^15,36^.

## Methods

### Samples

Germline samples (i.e. ejaculate or testis) and their corresponding soma samples (liver of the same individual or blood of the parents of that individual) are described in the **Supplementary Table 2**. In brief, ejaculate samples were collected from individuals of a domesticated zebra finch *T. g. castanotis* population (‘Seewiesen’) kept at the Max Planck Institute for Ornithology since 2004 (#18 in ^37^). Using a dummy female, we collected natural ejaculates from 8 brothers from two families (mitochondrial haplotype A, N_ejaculates_ = 9, from males A1-A3; mitochondrial haplotype B, N_ejaculates_ = 11, from males B2-B6). Males were 100-1036 days old (**Supplementary Table 2**). The study was carried out under license (permit no. 311.4‐si and 311.5‐gr, Landratsamt Starnberg, Germany). For details on the method of collection see the **Supplementary Methods**.

We collected testis samples and their corresponding soma from one captive *guttata* zebra finch (“Timor zebra finch”), which turned out to be a *castanotis* x *guttata* hybrid (see **Supplementary Material**), one wild-caught (male I, from Western Australia) and 11 captive *castanotis* zebra finches (“Australian zebra finch”). The *castanotis* samples were chosen to include nine major mitochondrial haplotypes (i.e. males A to I, A0, A1 and B1 from matrilines A to I; note that male A0 was an uncle of males A1-A3, and males B1-B6 were brothers; **Extended Data Figs. 1,2**; see **Supplementary Methods** for mitochondria sequencing and assembly). Matrilines A, B and E were first described in ^19^, whereas the others were described in this study and named as C, D and F to I for simplicity. The 11 captive *castanotis* zebra finches were sampled from two recently wild-derived populations (‘Bielefeld’ #19 in^37^ and ‘Melbourne’^38^) and three domesticated populations (‘Krakow’ #11 in^37^, ‘Seewiesen’ and ‘Spain’^8^). The sampled individuals were 238-1882 days old (**Supplementary Table 2**). All 11 captive *castanotis* zebra finch testes were large (longest diameter: 3-5 mm) compared to the testis of the *guttata* zebra finch (longest diameter: ~ 1mm), suggesting that the *guttata* male might have been sexually inactive and hence might have a lower fraction of germ cells within the testis compared to the *castanotis* samples.

### Cytogenetics

To determine the presence or absence of GRC in mature sperm, we conducted FISH on the ejaculates of one brother of matriline A and four brothers of matriline B (see sampling and **Supplementary Table 1**), following a protocol modified from^39^. In brief, we collected fresh ejaculate in 10 μl of PBS with a 20 μl pipette, and osmotic-shocked the sample by adding 250 μl (~ 20-fold volume of the sample) of 1% sodium-citrate solution for 20 minutes. We then spread the sample on a microscopy glass slide placed on a 60° C heating plate, following the Meredith’s technique^39^, and let it dry on the heating plate. We used a FISH probe^8^ for the gene *dph6*, which is in about 300 copies on the GRC but only represented as a single copy paralog on the A chromosomes^8^. We amplified the probe by PCR from DNA extracted from a *castanotis* zebra finch testis using the primer sequences F-ACGTCTTTGCCTGACCCTTTCAGA, R-TGCATAGAGTTCTCCATCAGACAGACA, taken from^8^, and then labelled it with tetramethylrhodamine-5-dUTP via nick translation^40^. FISH was then performed on the ejaculate preparations. The hybridization mix consisted of 12 μl formamide, 6 μl dextran sulfate, 1.5 μl 20×SSC, 0.5 μl salmon sperm, 0.5 μl SDS, 3 μl *dph6* probe, and 6.5 μl H_2_O. We applied 10 min of denaturation at 70° C.

FISH preparations were analyzed along the z-axis using a DSD2 confocal unit fitted to an Eclipse Ti inverted microscope (Nikon) with a Zyla sCMOS camera (Andor) under near-UV (405 nm) and green (550 nm) sequential excitation, obtained with a pE-4000 (CooLED) device, to determine whether the GRC signal from the GRC-carrying sperm was inside the nucleus. To quantify the fraction of GRC-carrying sperm in the ejaculate, we sampled 8 fields (2 rows x 4 columns) from each FISH preparation where there were more than 20 non-overlapping spermatozoa (hence, we did not use prior information on a GRC signal), using a Zeiss Axio Imager.M2 microscope equipped with DAPI (465 nm), red (572 nm) and green (519 nm) filters at 40x magnification. We took photos with a Zeiss Axiocam 512 colour camera using ZEN blue 3.1 software. We then counted the total number of spermatocytes, the number of GRC-carrying spermatocytes and the number of expelled free-floating GRC micronuclei for all sampled fields. Additional images were generated under Leica DM 6000/HX PC APO 100x-Oil immersion for visualization. Images were processed using Fiji^41^.

### Whole-genome sequencing

We extracted genomic DNA from testis, ejaculate, and liver samples using a phenol chloroform extraction (details see **Supplementary Methods**), and from blood using the NucleoSpin Blood QuickPure Kit (from the company Macherey-Nagel) following the manufacturer’s instructions. Details of library preparation methods and library size can be found in **Supplementary Table 2**. Note that the raw sequencing reads of *castanotis* males A (SR00100), B (Spain_1) and F (Spain_2) were taken from^8^.

To study individual repeatability of GRC elimination patterns and to compare between-family differences, we sequenced DNA from 15 ejaculates (see Samples) and from the blood of the four parents (founders of families A and B) and the sampled males (as somatic baseline) using PCR-free Illumina libraries (**Supplementary Table 2**). We also sequenced DNA from the testis of one son per family (A1 and B1 in **Fig. 2**) to compare the GRC content between the testis and the ejaculate, which provides a measure of the remaining GRC content in the ejaculate after elimination during spermatogenesis. Additionally, we sequenced DNA from testis and liver of an uncle of the individuals from matriline A (sample A0 in **Fig. 2**).

To study the genetic diversity of the GRC, we sequenced DNA from testes and liver samples of a single male from each of the two most common mitochondrial haplotypes from the domesticated ‘Krakow’ population, as well as from each of the most common mitochondrial haplotypes of the two recently wild-derived populations ‘Melbourne’ and ‘Bielefeld’.

All PCR-free libraries were constructed and sequenced with 20-fold coverage on the Illumina HiSeq 3000/4000 (ejaculate and blood samples) or the NovaSeq SP platforms (testis and liver; 2 × 150 bp, paired-end) at the Institute of Clinical Molecular Biology (IKMB) of Kiel University, Germany. Additionally, we generated 10X Chromium linked-read data for DNA samples that were extracted from testis and liver samples using magnetic beads on a Kingfisher robot (for details see Kinsella et al.^8^) from one captive *guttatta* x *castanotis* hybrid with *guttata* phenotype (see **Supplementary Results** and **Fig. 3**), one domesticated *castanotis* zebra finch (male E) from population ‘Seewiesen’ with a different mitochondrial haplotype (E *sensu*^19^) and one wild-caught *castanotis* zebra finch from Western Australia (male I). All Chromium libraries were constructed and sequenced to high coverage (for details, see **Supplementary Table 2**) as paired-end (2 × 150 bp) using the Illumina NovaSeq 6000 S4 platform at SciLifeLab Stockholm.

### Raw reads processing

Raw reads from all WGS libraries (PCR-free and Chromium) were processed following a modified version of the “Genome Analysis Toolkit Best Practices Pipeline”^42^. We filtered the raw reads using ‘BBDuk^43^’ and trimmed the last base of each sequence, putative adaptor sequences and bases with low quality, and only kept high-quality reads that were more than 50 bp long. We then mapped each library (paired-end reads) against the reference somatic genome, taeGut1^18^, using BWA-MEM v0.7.17 ^44^ with the default settings while marking shorter split hits as secondary. In a next step, we used the ‘Picard^45^ MarkDuplicates’ option to mark mapped reads that might result from PCR duplication, to reduce PCR-bias in the abundance of certain DNA fragments during sequencing. Finally, we analysed coverage and called SNPs (see below) for downstream analysis. For a detailed pipeline and the corresponding scripts see **Code accessibility**.

### Coverage analysis

We applied an analysis of sequencing coverage that was adapted from Kinsella et al.^8^. We calculated ratios of sequencing coverage of germline samples (ejaculate or testis) over somatic samples (liver or blood, averaged for the two parents when applicable) in non-overlapping adjacent windows of 1 kb width across the entire genome. For each library, we calculated read coverage using ‘SAMtools v1.6^46^ depth’ per bp, and used average values for each 1-kb window. For each germline sample, we then calculated the coverage ratio between germline and its corresponding soma library and log_2_-transformed the values, after correcting for variation in library size, i.e. by dividing the coverage-per-window by the total number of base pairs sequenced for that library.

To quantify testis enrichment in coverage (compared to soma), we first removed windows with too low coverage, i.e. those where both soma and germline samples had <3-fold coverage. Second, we calculated mean and standard deviation (SD) of coverage for all 1-kb windows of each somatic library and removed windows with coverage >2 SD above the mean of a given library. Such high coverage values indicate duplications on the A-chromosomal paralog, which makes quantification of copy number enrichment in testis difficult. Third, we centred the log_2_-transformed germline-to-soma coverage-ratios of the high-quality windows on the median of the above-selected windows.

To compare the amount of the GRC-linked DNA that remained in the ejaculates between the two focal families (matrilines A and B), we selected those 1-kb windows (N = 1,742) that showed significant testis coverage enrichment (i.e. log_2_ testis-to-soma coverage-ratio ≥ 2) using all individuals for which we sequenced testis DNA, but excluding those that were closely related to the males for which we sampled ejaculates (N = 9, see Samples and **Fig. 2a**). Then, we estimated the individual repeatability of the log_2_ ejaculate-to-soma coverage ratios of the selected windows (response variable) among the 15 ejaculate samples, using a mixed-effect model with the ‘lmer’ function in the ‘lme4^47^’ package in R v4.0.3^48^, in which we fitted individual, ejaculate, and window identity as random effects. Matriline repeatability was estimated by fitting matriline identity as an additional random effect in the previous mixed-effect model. To estimate the between-family difference in the GRC amount in ejaculates, we added ‘matriline’ as a fixed effect in the mixed-effect model of individual repeatability. To estimate the amount of reduction in the GRC content in ejaculate samples compared to the testis in each family (A and B), we used two linear models (one for each family; see **Fig. 2**), using the ‘lm’ function in the R package ‘stats’. Here we used the median of the log_2_ germline-to-soma coverage ratio (of the above selected windows) for each germline sample as response variable, because it is less sensitive to potential copy-number variation in the GRC-linked high copy number genes compared to the mean. In these two linear models, we only fitted the type of germline tissue (ejaculate or testis) as a fixed effect. For model structures and outputs, see also **Supplementary Table 3**.

### SNP analysis

We used the SAMtools v1.6^46^ mpileup tool to call SNPs, and a customized R script to filter for high-confidence SNPs of interest (see **Code accessibility**), as follows. To study the overall between-individual variation in GRC haplotypes, we called SNPs for all testis/soma pairs (one hybrid *castanotis* x *guttata* and 12 *castanotis* males) simultaneously and selected high-confidence sites (see **Supplementary Methods** for details). We then identified high-confidence testis-specific alleles by selecting sites for which (1) the soma library had more than 10 reads, (2) the allele was found in ≥ 3 reads in the germline sample, and (3) the allele was only present in the testis sample but not in the corresponding soma sample. We also identified those testis-specific SNPs that were private to only one of the 13 sequenced individuals, i.e. those testis-specific alleles that were absent from all soma and testis libraries except for the focal one. To reduce false positives in the private testis-specific SNPs that were identified above, we focused on regions that contained multiple private testis-specific SNPs, defined as 10-kb non-overlapping adjacent windows with at least three such private SNPs (**Extended Data Fig. 5 and Fig. 3b,c**).

To study the GRC content in ejaculates, we called SNPs simultaneously for mapped reads from the 15 ejaculates, the two testes of the brothers from A1 and B1 in **Fig. 2b,c**, the four blood samples of their parents and one pair of testis and liver of one uncle of family A (A0 in **Fig. 2b**). We filtered for high-quality, germline-specific alleles, following the same procedure as described above. We then identified the 1-kb non-overlapping adjacent windows that contained at least 10 germline-specific SNPs (**Extended Data Fig. 4**).

To study the extent of A-chromosomal introgression of *castanotis* DNA into the captive population of *castanotis* x *guttata* hybrids, we called SNPs for the combined soma libraries of the hybrid (liver) and a pool of 100 wild-caught *castanotis* zebra finches^27^ (blood). We then filtered for those high-quality SNPs that were homozygous in the (predominantly *guttata*) hybrid, but absent from the 100 *castanotis* zebra finches. Additionally, we filtered for SNPs that were heterozygous in the soma (liver) library of the hybrid individual. We calculated the number of fixed (i.e. homozygous) *guttata* SNPs and the number of heterozygous sites for non-overlapping adjacent windows of 500 kb. We considered windows with a low number of fixed *guttata*-specific SNPs as signals of *castanotis* introgression. We determined the copy number of those *castanotis-*introgressed sequences by their level of heterozygosity: a run of homozygosity would indicate two copies of one *castanotis* haplotype, a similar level of heterozygosity compared to the non-introgressed regions suggests two *castanotis* haplotypes, and an extremely elevated heterozygosity level implies that one copy of the *castanotis*-haplotype segregates with a *guttata* haplotype.

### Haplotype analysis

To estimate the genetic diversity among GRC haplotypes and the linkage disequilibrium between GRC haplotypes and mitochondrial haplotypes, we first generated a consensus for each GRC paralog, and then created haplotypes for each sample, as described below. Here, we focused on single-copy or low-copy-number GRC genes that are diverged from the A-chromosomal paralog. This is necessary to ensure a sufficient number of SNPs to bioinformatically differentiate the two paralogs (see section **Main**).

To generate a consensus of each GRC paralog, first, we used SAMtools v1.6^46^ view for each germline and soma library to subset reads that mapped on the A-chromosomal paralog (i.e. taeGut1^18^) of the focal gene. Second, we used MEGAHIT v1.2.6^24^ to assemble those mapped reads into contigs *de novo*. We then used MAFFT v7.429^25^ to align the assembled contigs that were longer than 2 kb. To build a consensus of the GRC paralogs, we manually selected the contigs that were absent from all soma libraries but present in more than half of the germline libraries, whereby we generated the consensus using the most common allele (for sites that contain SNPs). The consensus of *bicc1*_*GRC*_ and *pim3*_*GRC*_ were constructed from a single *de novo* assembled contig, while the consensus of the *elavl4*_*GRC*_ consisted of five *de novo* assembled contigs that were scaffolded with multi-N nucleotides in between them.

Using BWA-MEM v0.7.17^44^, we separated the reads from the GRC genes and their A-chromosomal paralogs by mapping reads against both the A-chromosomal (i.e. sequence on taeGut1^18^) and the GRC paralogs of that gene. For single-copy GRC genes, this allowed us to generate naturally-phased GRC haplotypes for each sample, and to check for heterozygosity in terms of GRC haplotypes. Then, we used SAMtools v1.6^46^ mpileup to call SNPs for each library of those mapped reads. For the two single-copy GRC genes, *bicc1*_*GRC*_ and *pim3*_*GRC*_ (**Fig. 4b,c**), we then generated one GRC haplotype for each germline sample by substituting the called alternative allele from the reference consensus allele using customized R scripts (see **Code accessibility**). For the double-copy GRC gene, *elavl4*_*GRC*_ (**Fig. 4d**), we artificially phased the reference allele of the consensus sequence into haplotype 1 and the alternative allele into haplotype 2. Unfortunately, the coverage of GRC-linked reads was too low for the hybrid male with the *guttata* GRC, so we were unable to construct the *guttata* version of these low-copy GRC-linked genes. Hence, we analysed the GRC-haplotypes of the *castanotis* zebra finches only.

To compare the genetic diversity of the GRC genes with their A-chromosomal paralogs, we used read aware phasing of SHAPEIT v2.r904^49^, to phase the two A-chromosomal haplotypes for each somatic library for the GRC genes (**Fig. 4**; see **Code accessibility**).

To compare the genetic diversity between different GRC genes and their A-chromosomal paralogs, we first used MAFFT v7.429^25^ to align the above-constructed haplotypes of GRC and A-chromosomal paralogs for each gene. We then used BMGE v1.12^50^ to trim positions with > 20% “missingness” (i.e. gaps between GRC and A-chromosomal paralogs) in each alignment. Finally, we used DnaSP v6.12.01^51^ to calculate the mean number of pairwise differences per site (π) for each GRC gene and each A-chromosomal paralog in the gap-free alignments, and used this value as an estimate of genetic diversity. For each set of genes (GRC and A-chromosomal paralogs), we analyzed and plotted a haplotype network using the ‘haplotype’ function from the R package pegas v0.14^26^ (see **Code accessibility**).

## Data accessibility

All NGS data will be deposited at the NCBI. All alignments will be deposited in figshare. Supporting data will be uploaded to the Open Science Framework.

## Code accessibility

Supporting scripts will be uploaded to the Open Science Framework.

## Authors’ contributions

Conceptualization: Y.P., W.F., F.J.R.R., A.S. and B.K.

Ejaculates sampling and sequencing design: Y.P., U.K. and W.F with inputs from J.C.M.

Testis and liver sampling and sequencing design: W.F., A.S. and Y.P.

Ejaculates collection: Y.P.

Tissue dissection: Y.P., K.T. and M.H.

DNA extraction: K.T.

PCR-free sequencing: A.F. and M.H.

Mitogenome amplicon sequencing design: Y.P. and W.F. with inputs from J.C.M and U.K.

Primer design: Y.P.

Mitogenome sequencing and de-multiplexing: S.B.

Sequencing data analysis: Y.P. with help from F.J.R.R. and inputs from W.F., A.S and J.C.M.

FISH: Y.P. with help from F.J.R.R. and inputs from J.C. and J.P.C.

FISH counting: I.G.

Confocal microscopy: J.P.C and J.A.

Cytogenetic interpretation: Y.P., W.F. with help from F.J.R.R., J.C., J.P.C. and M.H.

Results interpretation: Y.P., W.F. and A.S. with inputs from F.J.R.R.

Funding acquisition: B.K. and A.S.

Illustration: Y.P.

First draft of manuscript: Y.P.

Writing: Y.P., W.F., A.S. and B.K., with inputs from all authors.

## Acknowledgements

We thank Melanie Schneider and Christine Baumgartner for support with molecular work, Martin Irestedt for help with DNA extractions for the 10X samples, Shouwen Ma for discussion on microscopic image processing, and Katrin Martin, Isabel Schmelcher, Claudia Scheicher, Sonja Bauer, Edith Bodendorfer, Jane Didsbury, Annemarie Grötsch, Andrea Kortner, Petra Neubauer, Frances Weigel and Barbara Wörle for animal care and help with breeding zebra finches. We thank Leo Joseph and the Australian National Wildlife Collection for providing testis and liver samples from a wild *T. g. castanotis* individual. We thank Leo Joseph, Julie Blommaert, Octavio Palacios and Simone Fouché for comments on the manuscript. Some of the computations were performed on resources provided by the Swedish National Infrastructure for Computing (SNIC) through the Uppsala Multidisciplinary Center for Advanced Computational Science (UPPMAX). The authors acknowledge support from the National Genomics Infrastructure in Stockholm funded by Science for Life Laboratory, the Knut and Alice Wallenberg Foundation and the Swedish Research Council.

